# Generating snoRNA-guided Programmable 2’-*O*-methylation

**DOI:** 10.1101/2024.11.18.622494

**Authors:** Justin Zhao, Eric Cockman, Xinhe Yin, Yvonne Chao, Chad Pecot, Christopher L. Holley

## Abstract

Small nucleolar RNAs (snoRNAs) are critical in guiding post-transcriptional modifications like 2’-*O*-methylation (Nm), which play crucial roles in downstream processes such as splicing and translation. This study tests a novel method for Nm validation, addressing a significant gap in modern Nm research, and offers insight into the intricacies of snoRNA-guided Nm. While mapping of Nm modifications has seen significant improvement within the past decade, no major techniques have been able to validate these potential sites. Additionally, many mapping techniques lack consensus among proposed Nm sites, especially on mRNAs. Without a proper validation technique, Nm research lags compared to its peer post-transcriptional modifications. The RNase H-based Nm-VAQ assay used here quantifies 2’-*O*-methylation at single nucleotide resolution across various RNA species including rRNA, snRNA, and mRNA. Its optimization for mRNA allows for an unprecedented way to study the effects of Nm modifications in low abundance transcripts. Utilizing this, the study also explores the potential of creating synthetic snoRNAs to guide Nm modifications. Exogenous snoRNAs are shown to rescue Nm in genetic knockout models and can be mutated to guide Nm at any location along the target RNA transcript. Preliminary work indicates that synthetic snoRNAs demonstrate the ability to modify luciferase, impacting translation efficiency. Targeting an exon increases mRNA abundance but decreases protein expression, consistent with previous findings on *Pxdn* mRNA. These findings set the scene for novel understanding of the relationship between snoRNA abundance, 2’-*O*-methylation efficiency, and Nm’s impact on gene expression.

## Introduction

Small nucleolar RNAs (snoRNAs) are a class of highly expressed non-coding RNAs (ncRNAs) commonly found within eukaryotic organisms ranging from 60-300 nt^1^. They have been shown to play important roles in the post-transcriptional modification of rRNAs within the nucleolus, impacting rRNA folding, stability, and function as parts of the ribosome subunits^2^. SnoRNAs are commonly structurally characterized into three distinct categories: H/ACA box snoRNAs, C/D box snoRNAs, and small Cajal RNAs (scaRNAs). The former two are responsible for guiding pseudouridylation (Ψ) and methylation of their complementary RNA targets, respectively. ScaRNAs can guide either methylation, pseudouridylation, or both^3^. This function is accomplished by the association of the snoRNA with various protein subunits to form a small nucleolar ribonucleoprotein (snoRNP), which is then guided to the target by the snoRNA, and the modification is made^4^.

C/D box snoRNAs typically range from 60-90 nucleotides and contain two motifs: the C box (RUGAUGA, R=purine) and the D box (CUGA), which are located at the 5’ and 3’ ends of the transcript respectively. Many also usually contain less conserved duplicates of these two sequences, termed C’ and D’ boxes^5^. Base pairing between the 5’ and 3’ ends of C/D box snoRNAs causes the formation of a stem loop structure that is vital for proper assembly of the small nucleolar ribonucleoprotein (snoRNP)^6^. These snoRNAs associate with four separate proteins: fibrillarin (FBL), NOP56, NOP58, and 15.5K. The anti-sense element found upstream of the D or D’ boxes guides the snoRNP to its complementary region on the RNA target and fibrillarin, a methyltransferase, modifies the fifth nucleotide upstream of the D or D’ box by methylating the 2’-OH group of the ribose sugar^7,8^. This produces a post-transcriptional modification known as a 2’-*O*-methylation (Nm). This anti-sense element in each C/D box snoRNA is vital for guiding methylation to the correct nucleotide of its complementary RNA sequence. When these sequences are mutated, Nm modifications can theoretically be guided to an arbitrary rRNA sequence^9^. However, work in using synthetic snoRNAs to place modifications on various RNA species remains limited.

2’-*O*-methylation (Nm) is a post-transcriptional modification to RNA that attaches a methyl group (-CH_3_) to the 2’ hydroxyl (-OH) group of the ribose sugar in a nucleotide **(Figure 1)**. Nm can occur on any of the five RNA nitrogenous bases (A, C, G, U, and T) and is found on several types of RNA. They are highly abundant on rRNA, tRNA, and snRNA, allowing them to play roles in the ribosome, peptide chain formation, and the spliceosome^10–12^. These modifications have also been found with lower frequency on mRNAs, microRNAs (miRNAs), small-interfering RNAs (siRNAs), and Piwi-associated small RNAs (piRNAs)^13–16^.

**Figure 1.**
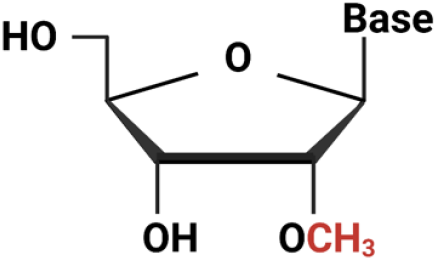
Example of a nucleotide with 2’-O-methylation (Nm) modification. The -H of the -OH group at the 2’ carbon of the ribose sugar has been replaced with a -CH3 group. This modification can occur on nucleotides with any of the 5 nitrogenous bases (A, T, U, C, G).

Due to their presence on mRNAs, Nm modifications play important roles in translation efficiency. The addition of an Nm inhibits translation, with the strongest inhibitory effect occurring when the modification is on the 2^nd^ nucleotide of the codon. This effect is conserved in both bacterial and eukaryotic systems, indicating its importance to proper organismal functioning^17,18^. The decrease in translation efficiency has been validated in a specific mRNA: *Pxdn*. Interestingly, while the presence of Nm modifications inhibited translation, it increased mRNA expression^19^.

Detection of Nm sites on RNA species, particularly lowly abundant transcripts such as mRNA, has remained a challenge in the field since the beginning of its study 70 years ago. Currently, four major techniques are in common use to map Nm modifications at single nucleotide resolution: RiboMeth-seq, 2’OMe-seq, RibOxi-seq, and Nm-seq^20–24^. While mapping of Nm modifications has seen significant improvement within the past decade, no major techniques have been proposed to validate these potential sites. As a result, there has been an accumulation of potential Nm sites with unknown stoichiometry and biological relevance, particularly in low abundance RNA species such as mRNAs. This issue is further compounded by the fact that among the four mapping techniques listed above, there is a very limited overlap of proposed sites, suggesting that the accuracy and precision of these methods can be improved. In comparison, other RNA modifications such as *N*^6^ – methyladenosine (m^6^A) have seen vast growth in study over the past two decades owing to the development of high-throughput mapping and validation techniques^25^.

One notable validation technique for 2’-*O*-methylation is reverse transcription at low dNTP concentrations followed by polymerase chain reaction (RTL-P). Reverse transcription is performed under two conditions: the first at high dNTP concentration (40 μM – 1 mM) and the second at low dNTP concentration (0.5 μM – 4 μM). In the case of low abundance transcripts such as mRNA species, a gene specific RT primer can be used instead of random hexamers or oligo dT^26^. Following this with qPCR allows for a semi-quantitative validation of Nm modifications, although it is not single nucleotide resolution. Despite these shortcomings, this method has proven useful in determining snoRNA-guided 2’-*O*-methylation even in mRNAs, as evidenced by the validation of an Nm modification in *Peroxidasin (Pxdn)*^19^. While RTL-P has potential to assist in the validation of Nm modification sites, there is still the need for a fully quantitative, single nucleotide resolution validation technique if the field is to advance at the same rate as other RNA modifications.

Ribosomal protein L13a (RPL13A) is a part of the L13 family of proteins and is a component in the 60S ribosomal subunit^27^. In addition to encoding for the RPL13A peptide, the gene also transcribes four intronic box C/D snoRNAs: *U32A, U33, U34*, and *U35A*. As box C/D snoRNAs, these four transcripts guide FBL-dependent 2’-*O*-methylation using their anti-sense elements. Although they are not 100% canonical C/D box snoRNAs (the anti-sense element is not directly upstream of the D box in *U34* and *U35A*), the anti-sense elements themselves map directly to sites found on 28S and 18S rRNA. Interestingly, *U32A* has two separate anti-sense elements, one directly upstream of the D box and the other directly upstream of the D’ box. It is predicted that the one upstream of the D box guides 2’-*O*-methylation of G1328 in 18S and the other guides methylation of A1511 in 28S. *U33* is predicted to guide U1326 in 18S, *U34* guides U2824 in 28S, and *U35A* guides C4506 in 28S^28^. These Nm sites on rRNA play important roles in the development of validation techniques as they are well-characterized with known methylation levels.

In an effort to advance the 2’-*O*-methylation field, the goal of the project was two-fold. First, a validation technique needed to be developed that could effectively measure whether an Nm modification was present at the specified site and the degree of methylation. Techniques have been proposed in the past but require further development and optimization to ensure results are consistent and replicable. Since Nm is well characterized on rRNA already, it was important to ensure the assay functions for low abundance mRNAs as well. Second, we wanted to determine the feasibility of synthetically adding a modification on an mRNA transcript using programmable snoRNAs, where the anti-sense element upstream of the D and D’ box was mutated to match the target sequence. This supported the creation of a model system for improving our understanding of the biological effects of Nm modifications at various sites on mRNA. Through the model, we were able to demonstrate that Nm affects downstream translation as well as transcription when occurring in an exon, consistent with previous findings in *Pxdn* mRNA. This work establishes a flexible system for future investigation of how snoRNAs guide Nm modification at additional sites on mRNA and pre-mRNA, such as the 5’- and 3’-UTR, start and stop sites, splice donor and acceptor sites, and introns.

## Results

### Optimization of RNase H protocol (Nm-VAQ) to quantitatively measure Nm modification

RNase H recognizes and binds to RNA-DNA duplexes, cleaving the nucleotide at the 3’-end or one nucleotide downstream **(Figure 2a)**^29^. This property is used to generate an RNA-DNA chimera that binds to the desired site and allows for RNase H to act. However, if an Nm modification is present at either of these sites, RNase H cleavage is blocked. The RNA can then be converted to cDNA and quantified through RT-qPCR. The cleavage efficiency, which indicates presence of methylation, can then be calculated by comparing between the samples incubated with and without RNase H. A schematic of the Nm-VAQ assay is shown **(Figure 2a)**^29^. On 18S rRNA of 293T WT cells, the A159 site is highly methylated; therefore, the majority of RNase H cleavage at A159 is blocked due to the modification. Since cleavage is blocked, there should be a high calculated percent methylation. To validate the methylation results from the RNase H protocol adapted from Tang et al., a control chimera was designed for the A159 Nm site at U161 that should not be cleaved^29^. U161 is two nucleotides away from Am159 and is known to not be methylated based on mass spectrometry analyses, so RNase H activity should not be hindered. Full cleavage of the chimera-RNA complex should occur at this site, resulting in a low calculated percent methylation. In order to decrease stochastic RNA cleavage, 10 μM Tris pH 7.0 buffer was used in the initial annealing reaction instead of nuclease-free water, and the reaction was moved immediately onto ice after incubation to prevent base-catalyzed hydrolysis of RNA that disrupts the assay. All reactions were gently mixed by pipette instead of vortex to prevent disruption of the RNA/chimera complex after annealing and ensure that complete cleavage occurs following incubation with RNase H. The two steps of dilution are necessary to dilute out the RNase H enzyme and excess chimera that interfere with cDNA synthesis.

**Figure 2.**
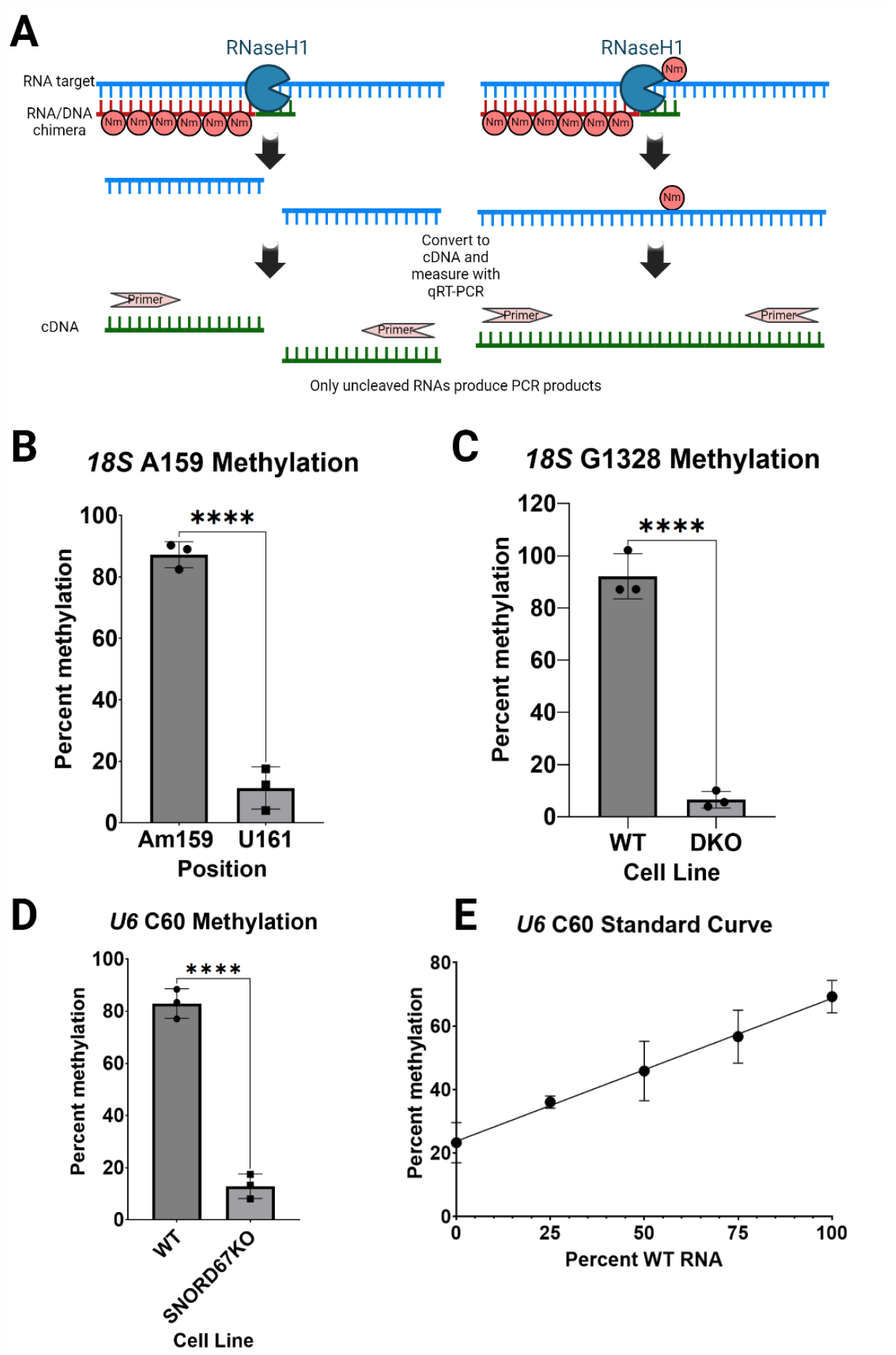
Establishment and optimization of Nm-VAQ. Nm quantification on the A159 and G1328 sites of*18S* rRNA were performed in HEK293T cells and D3H2 cells for the *U6* snRNA C60 site. **A)** Schematic of RNase H cleavage and detection. RNase H cleaves sites that do not contain an Nm modification. Illustration created with BioRender.com. **B)** The control chimera demonstrates the robustness of Nm-VAQ at single nucleotide resolution. Compared with non-methylated U161: ****, p < 0.0001. **C)** Nm-VAQ’s single nucleotide specificity can be recapitulated in a genetic knockout model. Compared with *U32A/U51* DKO: ****, p < 0.0001. **D)** Nm-VAQ can be applied to a less abundant snRNA species. Compared with SNORD67KO: ****, p < 0.0001. **E)** Nm-VAQ demonstrates linearity across a standard curve when tested on *U6* snRNA. Linear regression: R^2^ = 0.89.

The modified Nm-VAQ assay performed on the 18S rRNA A159 site and U161 control site showed 87% methylation for the A159 site, consistent with previous literature on the site and other rRNA Nm sites^30^. Meanwhile, the control site showed 11% methylation, indicating that the modified Nm-VAQ assay can quantify Nm status at single nucleotide resolution but that there is some background signal at this nearby unmethylated site **(Figure 2b)**. To further test the robustness of the assay, Nm-VAQ was performed on the 18S rRNA G1328 Nm site with the 293T and 293T *SNORD32A* and *SNORD51* double knockout (DKO) cell lines. The G1328 Nm modification is guided by *SNORD32A* and is therefore absent in the DKO cells. The G1328 site showed 92% methylation in WT cells and 6.5% methylation in the DKO cells, further demonstrating that the optimized Nm-VAQ protocol can quantify methylation status at the single nucleotide level, but again that unmethylated sites have some level of background signal **(Figure 2c)**.

After demonstrating that Nm-VAQ could accurately quantify methylation at specific sites, the next goal was to apply the assay to RNAs of lower abundance such as snRNAs. The C60 methylation site on snRNA *U6* was targeted for this purpose. These experiments were conducted in the MDA-MB-231-D3H2LN (D3H2LN) human breast adenocarcinoma cell line. The Cm60 on *U6* is guided by *SNORD67*, so a *SNORD67* knockout line in D3H2 cells was used as a negative control instead of a separate control chimera targeting a nearby site. The C60 site showed 83% methylation in WT D3H2 cells and 13% methylation in the D3H2 *SNORD67* KO cells **(Figure 2d)**. A mixing experiment was also done by mixing proportions of WT and *SNORD67* KO cells to determine if Nm-VAQ could demonstrate linearity. Mixes of 0%, 25%, 75%, and 100% D3H2 WT RNA were used and Nm-VAQ was performed. The calculated methylation percentages demonstrated linearity across a range of 24% methylation for the 0% WT RNA to 69% methylation for the 100% WT RNA samples **(Figure 2e)**. Interestingly, a 0% methylation status was never achieved, even in the genetic knockout models. This alludes to the high background noise present in these experiments, which can be attributed to incomplete cleavage at unmethylated sites, which could be due to RNA secondary structure or other RNA-RNA interactions that do not lead to RNase H-mediated cleavage. Methylation status was also never measured at a full 100%; instead, values tended to hover around the 80-90% methylation range. This was likely due to unavoidable RNA degradation during the heating portions of the Nm-VAQ assay, although it was reduced with the usage of Tris pH 7.0 buffer instead of water. Even with these shortcomings, these experiments demonstrated that Nm-VAQ could be applied across cell lines and different types of RNA to quantitatively determine methylation status at single nucleotide resolution.

While Nm-VAQ demonstrated robust results with RNA species occupying varying magnitudes of abundance, the detection of cleavage among some of the lowest abundance transcripts, mRNAs, remained a challenge. To combat this, total RNA was poly-A selected to enrich for mRNAs that would then be used in Nm-VAQ. To further improve qPCR detection after cleavage with RNase H, the dilution steps were removed and replaced with a phenol:chloroform:isoamyl alcohol extraction followed by column cleanup. Simply removing the dilution steps was insufficient as the presence of residual RNase H enzyme and buffer inhibited downstream cDNA synthesis. The addition of this extraction step inactivated the RNase H enzyme without the need for a 90 °C heating step after cleavage as well, further protecting the RNA from base-catalyzed hydrolysis which would skew the results.

*PLXNB2, RPL10*, and *TOP1* were identified as potential candidates for mRNAs with Nm modifications based on various mapping techniques. G498 for *PLXNB2*, G150 for *RPL10*, and U877 for *TOP1* were the potential sites of 2’-*O*-methylation^29^. To combat false positives, control chimeras were designed to target G500, G152, and U879, respectively, for the three transcripts. All three control chimeras demonstrated near-complete cleavage of their respective sites with low variance, demonstrating that the optimized Nm-VAQ assay for mRNAs was successful. The percent methylation at the control chimera site (background signal) for *RPL10* was 23% **(Figure 3a)**, 4.5% for *TOP1* **(Figure 3b)**, and 17% for *PLXNB2* **(Figure 3c)**. Interestingly, only *PLXNB2* out of the three tested mRNAs with potential Nm modifications returned a positive result. *PLXNB2* demonstrated 74% methylation at the mapped site **(Figure 3c)**, while *RPL10* had 18% methylation **(Figure 3a)** and *TOP1* had 8.7% methylation **(Figure 3b)**. Neither *RPL10* nor *TOP1* had a significant change between the experimental and control chimeras, indicating there was likely no methylation at that mapped site. However, there was a significant difference between the two chimeras for *PLXNB2*, suggesting that the site was correctly mapped. The results from optimization of the Nm-VAQ protocol establish a novel validation method for any potential Nm modified sites on RNA species at differing magnitudes of abundance.

**Figure 3.**
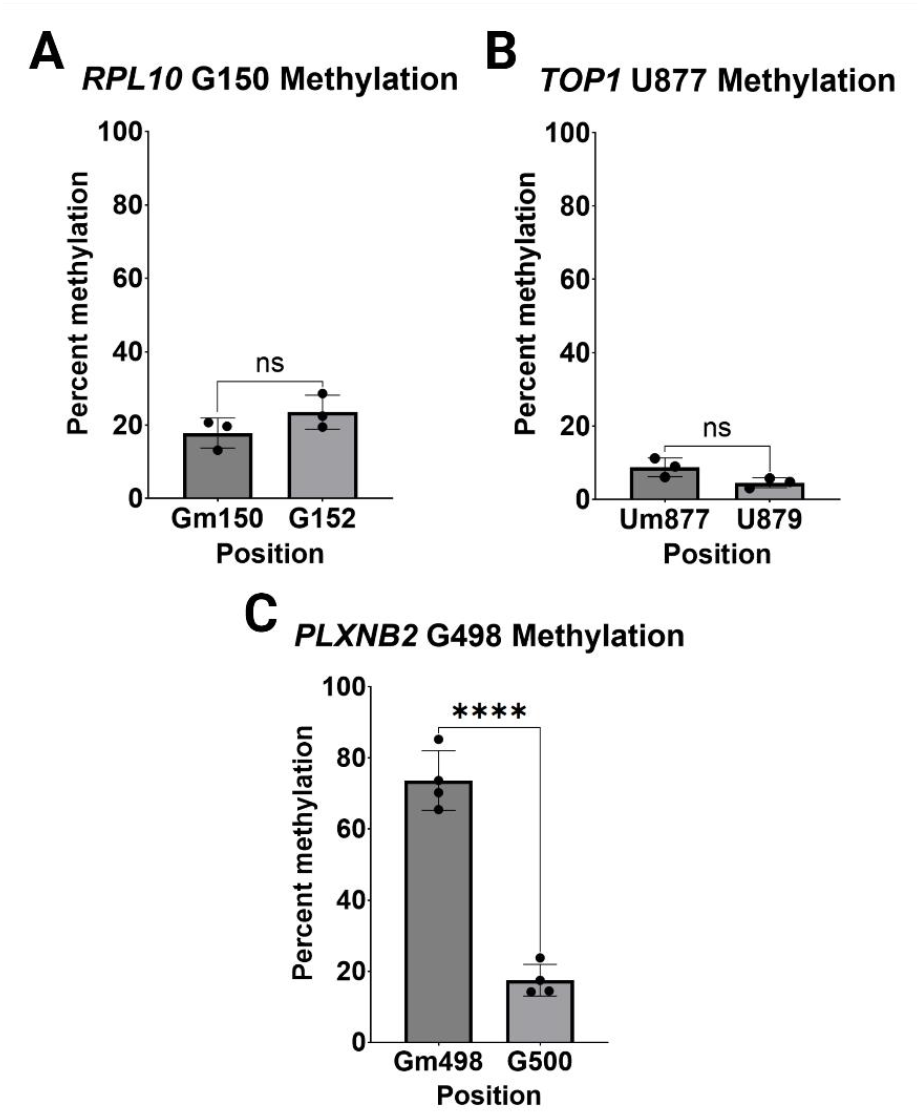
Optimization of Nm-VAQ for mRNA. **A)** *RPL10* shows no methylation at the proposed Gm150 site. Compared with G152: ns, p = 0.1882. **B)** *TOP1* also demonstrates no methylation at the Um877 site despite being previously mapped. Compared with U879: ns, p = 0.0649. **C)** *PLXNB2* Gm498, mapped by Tang et al., resulted in 74% methylation. This indicates a 2’-*O*-methylation exists at the Gm498 site. Compared with G500: ****, p < 0.0001.

### SNORD32A abundance increases following reverse transfection

Having established a method for quantifying 2’-*O*-methylation status at single nucleotide resolution, the next goal was to determine the effects of Nm modifications, specifically on less studied RNA species such as mRNAs. In order to accomplish this, we sought to synthetically add an Nm modification to certain mRNA transcripts and measure the effects on transcription, translation, and alternative splicing. This could then be correlated with the percent methylation of each transcript. As a proof-of-concept experiment to demonstrate that synthetic snoRNAs can methylate endogenous RNA, we *in vitro* transcribed (IVT) *SNORD32A (U32A)* and transfected it into a HEK293T cell line where *SNORD32A* and *SNORD51* were knocked out (HEK293T DKO). The *SNORD32A* oligonucleotide was reverse transfected into DKO cells to check whether non-native snoRNAs can act as replacement guides for snoRNA-guided Nm modifications. *In vitro* transcribed *SNORD32A* was first run on a 1% agarose gel as a quality control check to ensure the sample was not degraded or contaminated (not shown). 48 hours post-transfection with synthetic *SNORD32A*, RNA was extracted twice with Trizol reagent and run on a 1% agarose gel for quality control (not shown).

In order to demonstrate that transfection of exogenous snoRNAs could rescue methylation, we first had to show that reverse transfection conditions were conducive for *SNORD32A* to enter the cell. Therefore, *SNORD32A* abundance was measured across all transfection samples. Compared to non-transfected 293T WT cells, non-transfected DKO cells demonstrated essentially no *SNORD32A* expression. The transfected WT and DKO samples demonstrated similar levels of *SNORD32A* abundance, approximately 13-fold for both, compared to the non-transfected WT sample **(Figure 4)**. This indicates that 48-hour reverse transfection conditions are able to recover *SNORD32A* past native levels, identifying an avenue for recovering 2’-*O*-methylation in samples where the modification has been ablated.

**Figure 4.**
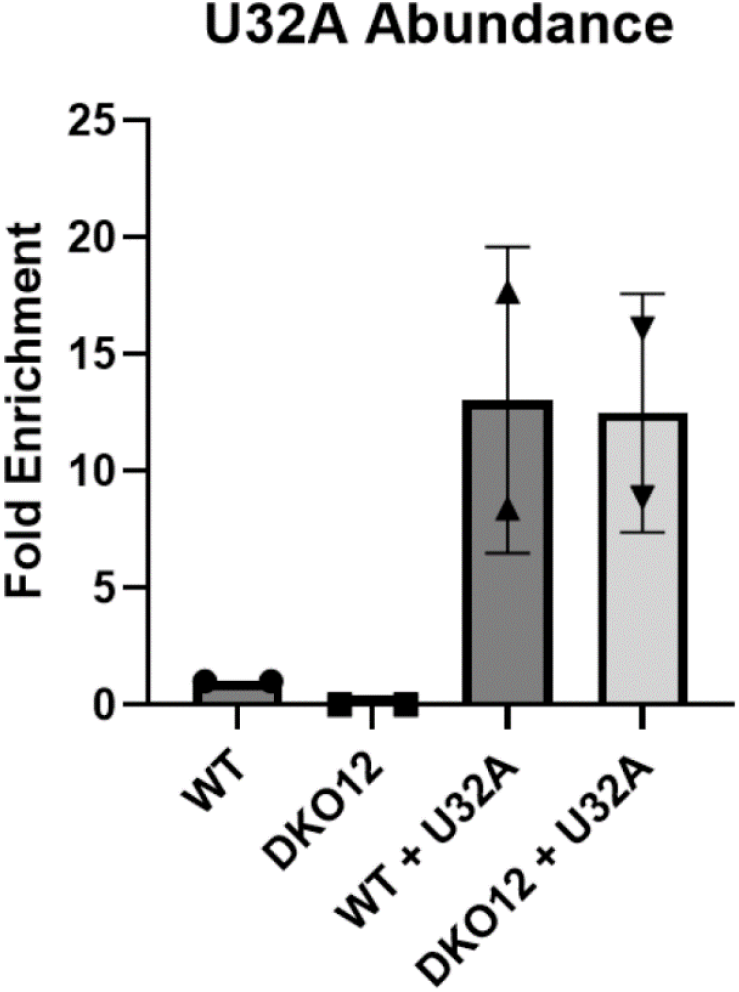
*SNORD32A* abundance following reverse transfection. *SNORD32A* abundance is increased past native levels following reverse transfection of synthetic snoRNA. Following the add-back of *U32A*, both WT and the *U32A/U51* DKO exhibited 13-fold enrichment of *U32A*. Abundance is essentially zero in DKO cells, indicating complete knockout of native *SNORD32A*.

### Transfection of SNORD32A rescues rRNA methylation

The anti-sense elements of *SNORD32A* and *SNORD51* target the G1328 site on 18S rRNA. We theorized that transfection of *SNORD32A* would rescue Nm modifications in *SNORD32A/51* DKO cells. As such, we used the newly optimized Nm-VAQ protocol to test for methylation status at this site following a 48-hour reverse transfection. The assay was performed on both the transfected samples and non-transfected negative control samples. Non-transfected WT cells showed 72% methylation at the G1328 site, and non-transfected DKO cells had 13% methylation, consistent with prior results shown in **Figure 2c**. *U32A*-transfected DKO cells demonstrated a rescue of methylation at the G1328 site, with 36% methylation, an approximately 3-fold increase from the non-transfected DKO cells **(Figure 5)**. Interestingly, *U32A*-transfected WT cells had 76% methylation at the G1328 site, indicating that overexpression of *SNORD32A* did not significantly increase methylation past native levels.

**Figure 5.**
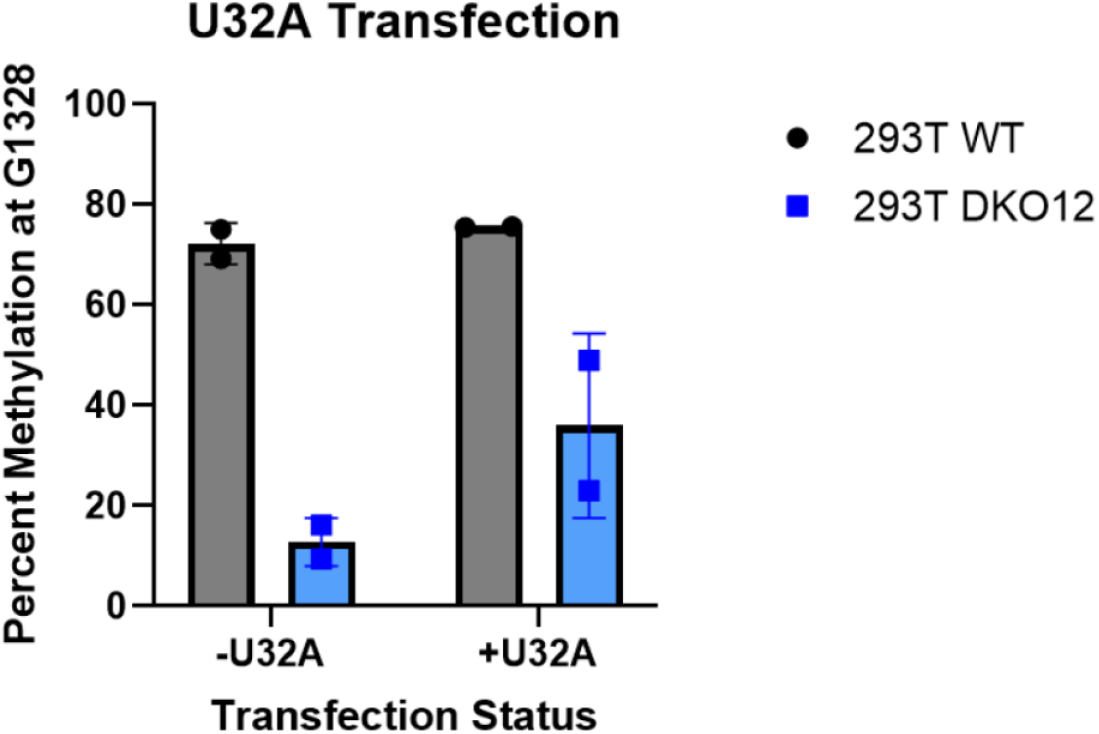
Transfection of synthetic *SNORD32A* rescues methylation at 18S G1328. HEK293T WT or DKO cells were transfected with synthetic *SNORD32A* and Nm-VAQ was performed to measure methylation status at the 18S rRNA G1328 site. The addition of exogenous *SNORD32A (U32A)*, successfully rescues methylation from 13% to 36% at the Gm1328 site on *18S* rRNA.

### 2’-O-methylation in exons decreases translation efficiency and increases transcript abundance

We then hypothesized that the effects of snoRNA-guided 2’-*O*-methylations on transcription, translation, and splicing could be investigated by targeting *Luciferase (Luc)* mRNA (or pre-mRNA) in stably transfected cell lines. Synthetic snoRNAs were generated using the backbone of an endogenous snoRNA, *U34*, but with an engineered antisense element that aligned to a site on the *Luc* transcript rather than its usual rRNA target. Using this method of mutating the anti-sense element, Nm modifications could be directed to various locations along the *Luc* transcript. Luciferase assays and RNA expression were then tested after transfection with these synthetic snoRNAs to determine the effects of the modification. A schematic of the model is shown below **(Figure 6)**.

**Figure 6.**
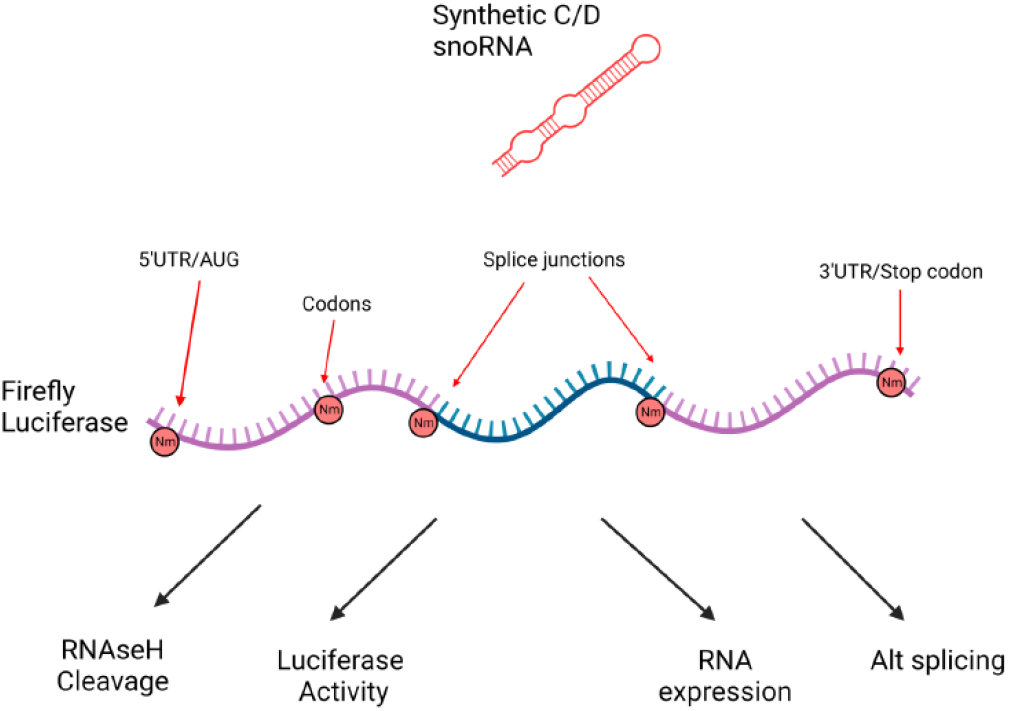
Schematic of the Luciferase experimental system. Possible targets for synthetic snoRNA-guided methylation on the *Luciferase* transcript include the 5’UTR/3’UTR, exons, splice junctions, and start/stop codons. Synthetic snoRNAs can be generated using the backbone of an endogenous snoRNA and mutating the anti-sense element. Nm can be measured using Nm-VAQ and downstream effects can be measured using Luciferase activity and qRT-PCR. Illustration created with BioRender.com.

Preliminary experiments for this Luciferase model system involved targeting a snoRNA to place an Nm modification to the second nucleotide of the Lys130 codon in the *Luc* transcript. This site was chosen based on previous work demonstrating that methylations at the second nucleotide of a codon introduced significant translational stalling^17,18^. Further work on *Pxdn* mRNA has demonstrated that Nm modifications on a lysine codon decreased translation efficiency and increased transcript abundance^19^. A negative control engineered snoRNA was generated by further mutating the methylation site 5 nucleotides upstream of the D box. We mutated the original nucleotide at the target methylation site from a U to a C in the control. Since box C/D snoRNAs have canonical Watson-Crick base pairing with their target RNAs, we theorized that mutation of the methylation site should impair snoRNA-guided Nm. In fact, previous studies have demonstrated that mutations at the target site disrupt methylation of the target RNA^19,31^. The change in translation efficiency was measured by proxy using the luciferase assay. The 0.2 nM snoRNA transfection conditions produced negligible change in the amount of luciferase activity, but at the 2 nM condition, the Lys130 snoRNA decreased luciferase activity to 49.52% of the 0 nM condition and the negative control decreased to 48.03% **(Figure 7a)**. Interestingly, the mutant synthetic snoRNA that should not be guiding an Nm modification to the *Luc* transcript caused similar significant decreases in protein expression as the experimental snoRNA. RT-qPCR was used to compare the levels of *Luc* mRNA in the 0.2 nM and 2 nM transfection conditions to the 0 nM condition. At 0.2 nM, both the Lys130 and Lys130 mutant snoRNA caused insignificant changes in mRNA expression. However, at 2 nM, both snoRNAs caused a significant increase in mRNA expression, with 1.7 and 1.9 times the expression compared to the 0 nM condition when normalized to GAPDH **(Figure 7b)**.

**Figure 7.**
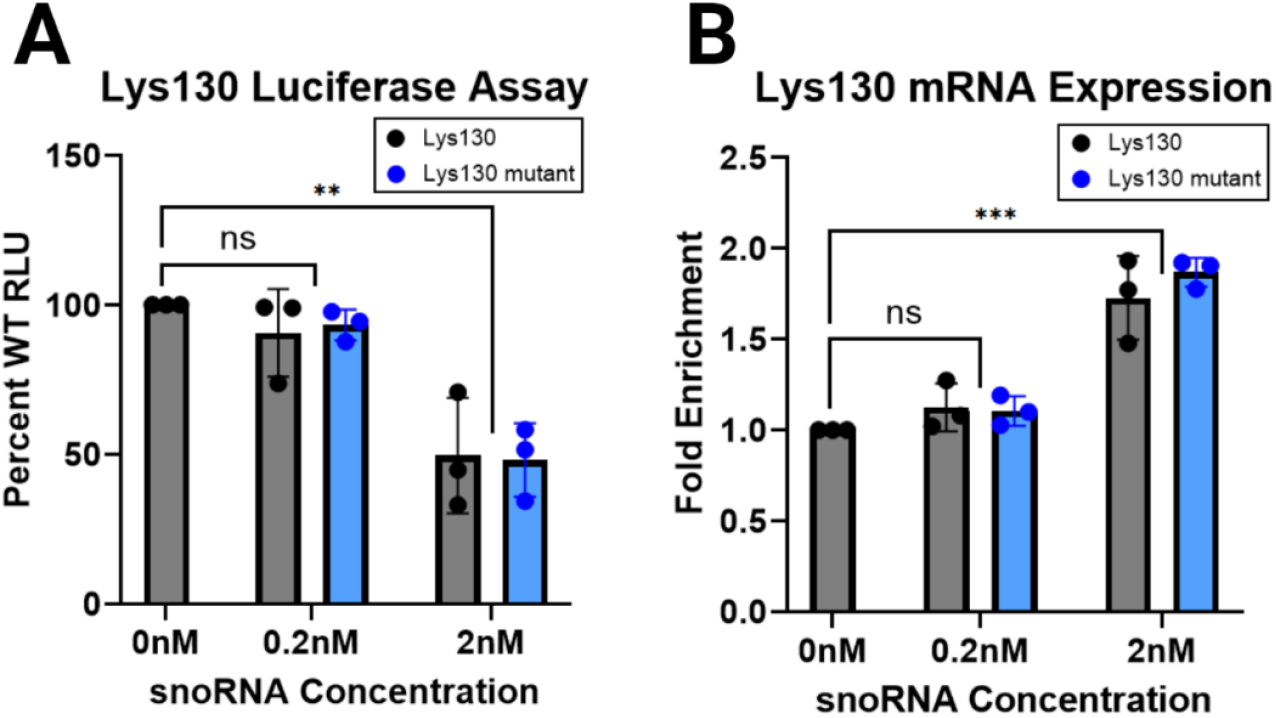
Functional changes in Luciferase expression when targeted by an engineered box C/D snoRNA. **A)** Addition of 0.2nM snoRNA did not reveal significant changes in Luciferase activity. Compared with %WT relative light units (RLU): ns, p = 0.66. At the 2nM condition, addition of the Lys130 snoRNA or a mutant that should not guide methylation reduced protein expression by 50%. Compared with %WT RLU: **, p = 0.0036. **B)** For the 0.2nM snoRNA transfection condition, there were no significant changes in mRNA expression, consistent with the effect observed in protein activity. Compared with WT: ns, p = 0.70. In the 2nM condition, both the experimental and control synthetic snoRNAs increased mRNA expression to 1.7- and 1.9-fold enrichment over the 0nM control respectively. Compared with WT: ***, p = 0.0007.

## Discussion

The Nm-VAQ protocol adapted from Tang et al. was described without a relevant negative control and an inability to determine if the percent methylation quantification was due to methylation presence or noise^29^. Here, the use of a control chimera that targeted a non-methylated site two nucleotides upstream from the target site allowed for validation of the Nm-VAQ assay. Addition of the control chimera revealed pitfalls in the original Nm-VAQ protocol of inconsistent cleavage that meant the low levels of methylation described in previous reports may be unreliable. Incomplete cleavage may be due to a variety of reasons, namely base-catalyzed hydrolysis of RNA, disruption of the RNA-chimera complex due to vigorous mixing, and inhibition of cDNA synthesis due to presence of RNase H enzyme. These three problems were solved through the optimization process: Tris 7.0 pH buffer was added to minimize RNA hydrolysis and the immediate transfer of the annealed reaction to ice instead of gradual cooling using the thermal cycler served to slow down the kinematics of hydrolysis. Mixing by pipette instead of vortex minimized disruption of the annealed RNA-chimera complex to ensure the RNase H was appropriately guided to the cleavage site. Additionally, the 1:50 dilution was necessary to reduce the impact of RNase H on cDNA synthesis and the following RT-qPCR. Without the dilution, presence of RNase H inhibited cDNA synthesis and caused a false positive methylation signal from Nm-VAQ. Optimization of the assay has established a robust tool for quantification of Nm.

Applying Nm-VAQ to previously mapped potential modification sites on mRNAs, such as *TOP1* and *RPL10*, revealed a distinct lack of methylation at the n or n+1 nucleotides. However, we were able to successfully validate an Nm modification on the *PLXNB2* mRNA, demonstrating the utility of Nm-VAQ in confirming putative sites identified through various transcriptome-wide mapping approaches. Notably, the negative results concerning *TOP1* and *RPL10* are not indicative of complete misses; since Nm-VAQ detects methylation at single nucleotide resolution, being just a couple nucleotides off during the mapping process would result in no methylation detection. It is entirely possible that an Nm modification could be present in the general region of the aforementioned mRNA transcripts, just not at the specific mapped nucleotide. It is in this instance where RTL-P can play a more effective role than Nm-VAQ since RTL-P validates whether a methylation exists in the larger region rather than a single nucleotide. However, one important issue exists where it is difficult to amplify and detect low abundance transcripts during RT-qPCR. This is especially prevalent for RTL-P where the low dNTP conditions result in significantly lower cDNA amounts as opposed to normal dNTP conditions.

This study also demonstrated the feasibility of using exogenous snoRNAs to rescue Nm modifications in a genetic knockout model. By transfecting *in-vitro* transcribed *SNORD32A* into a *SNORD32A/SNORD51* double knockout cell line, we rescued methylation at a well-characterized site on 18S rRNA that is guided by *SNORD32A*. Although we were able to transfect cells with greater than 10 times the amount of *SNORD32A* in a WT cell, methylation was not fully rescued at the canonical rRNA site. This could be due to the long half-life of rRNA, and a 48-hour transfection was not adequate for a large amount of rRNA turnover to occur. Future experiments should be undertaken to determine the ideal conditions for complete rescue of this methylation in a knockout cell line. With this proof-of-concept experiment that utilized our newly developed Nm-VAQ protocol, though, we demonstrated that Nm modifications could successfully be added to a HEK293T cell line using reverse transfection conditions. This paved the way for the study of generating targeted Nm modifications at specific nucleotides of a transcript.

To investigate the functional effects of Nm modifications on mRNA and protein biology, we established a model system using a stably transfected Luciferase reporter. In order to determine the effects of 2’-*O*-methylation on transcription and translation, an mRNA had to be expressed with no opportunities to be methylated by any endogenous snoRNAs. Eukaryotic firefly luciferase (Luc) was chosen due to its absence in the human genome and its capacity as a translational reporter gene. As such, the *Luc* mRNA transcript should only be modified by synthetic snoRNAs, and the effect of Nm modifications on translation efficiency can also be measured through a luciferase assay. The generation of the Luciferase expressing cell lines and synthetic snoRNAs marks the creation of a robust tool for investigating the effects of Nm modifications.

A notable finding was that targeting 2’-*O*-methylation at the second position of a lysine codon markedly decreased translation efficiency, which was measured by proxy using Luciferase protein expression. This decrease in protein was accompanied by an increase in mRNA expression, consistent with previous findings done on *Pxdn* mRNA. Interestingly, this change was observed for both a snoRNA that targets modification at that position, as well as a negative control that was mutated to prevent modification. One possibility for this positive result is that a single nucleotide mutation at the theorized modification site is not enough to completely prevent methylation.

Further work mutating various amounts of the anti-sense element can be performed to test whether this is truly the case. Another possible explanation involves the snoRNA-mRNA duplex that is formed when the snoRNP complex binds to the *Luciferase* transcript, where a stable RNA-RNA interaction (and recruitment of snoRNP proteins) could possibly inhibit translation without guiding Nm modification. Therefore, translational inhibition would occur despite any methylation being found on the transcript. A critical next step that is beyond the scope of this current work is to apply the newly optimized Nm-VAQ assay to measure Nm modification percentage at the Lys130 site to directly measure whether these synthetic programmable snoRNAs are truly guiding methylation at the site. Furthermore, other sites on the *Luc* mRNA will be tested such as the start and stop codons, UTRs, and introns to better elucidate the effect of Nm on transcription, translation and alternative splicing. Overall, these findings suggest that Nm modifications on mRNA can modulate both transcription and translation stability, playing important roles in broader gene expression.

In summary, this study provides a powerful tool for validating and quantifying 2’-*O*-methylation sites, addressing a significant gap in the field that has slowed down its development. We have additionally demonstrated the ability to guide methylations using exogenous snoRNAs to rescue methylation at canonical ribosomal Nm sites. Intriguingly, our results highlight the potential of synthetic snoRNAs to guide the placement of Nm to novel sites in the transcriptome. Lastly, our work has uncovered important roles that C/D box snoRNAs and Nm modifications play in mRNA and protein expression, influencing our understanding on the intricacies of epitranscriptomic gene regulation.

## Methods

### Cell lines

293T and other modified 293T cell lines were maintained in Dulbecco’s Modified Eagle Medium (DMEM) containing 10% Fetal Bovine Serum (FBS) from passages 10-24. Details on the modified 239T cell lines can be found in Elliot et al^19^. 293T cells were transfected with pairs of sgRNAs targeting *SNORD32A* and *SNORD51* using TransIT-LT1 (Mirus). D3H2LN cell lines were derived from Jenkins et al^32^. Details on the generation of the *SNORD67KO* D3H2LN cell lines can be found at Chao et al^33^. Briefly, cells were transfected with CRISPR/Cas9 constructs targeting *SNORD67* through electroporation. D3H2LN and *SNORD67KO* D3H2LN cell lines were maintained in RPMI containing 10% FBS.

### RNA and RT-qPCR

Total RNA was isolated from cells using Trizol (#15596026). RT-qPCR was run using PowerSybr (#4367659) and a StepOnePlus (ABI) Real-Time PCR System.

### Synthetic snoRNA reverse transfection

Synthetic snoRNAs were synthesized using the T7 RiboMax Express Large Scale RNA Production Kit (#P1320) and re-suspended in 30 μL of sterile nuclease-free water. The sequences are listed in the Supplementary Methods. This synthetic snoRNA was complexed with 1.5 μL Lipofectamine RNAiMax transfection reagent (#13778150) and 200 μL Opti-MEM (#31985070) for a concentration of 2 nM synthetic snoRNA. This concentration of snoRNA was chosen based on previous work in the lab. Cells were trypsinized and resuspended in 10 mL of DMEM containing 10% FBS. 293T cells were diluted with 1 mL cells in 5 mL DMEM while 293T DKO cells were diluted with 2 mL cells in 5 mL DMEM. 200 μL of the snoRNA-Lipofectamine RNAiMax complex was mixed with 800 μL of the cell dilutions in a 6-well plate. 200 μL of the snoRNA-Lipofectamine RNAiMax complex was mixed with 300 μL of the cell dilutions in a 12-well plate for all luciferase experiments. 200 μL transfection reagent mixture without any synthetic snoRNA was added to negative controls instead. The final volume was brought up to 2 mL for all 6-well plates and 1 mL for all 12-well plates with DMEM containing 10% FBS. Reverse transfections were extracted 48 hours post transfection.

### RNase H Cleavage

2’-*O*-methylated RNA/DNA chimeras targeting the methylation site were synthesized through Dharmacon Horizon. Chimera sequences are listed at the end of the Methods section. 500 ng of total RNA from 293T cells was mixed with 50 pmol of the RNA/DNA chimera and brought up to a total of 11 μL with 10 μM Tris pH 7.0 buffer and mixed thoroughly by pipette. For mRNAs specifically, 500 ng of polyA-selected RNA was used instead of total RNA. The samples were incubated at 95°C for 1 minute and transferred to ice immediately. 5 μL of the annealed RNA and chimera was mixed with 1 μL of RNase H enzyme (#M0297S) and 1 μL of 10x RNase H buffer (#B0297S) and the final volume was brought up to 10 μL with nuclease-free water. The other 5 μL of the annealed RNA and chimera was mixed with 1 μL of 10x RNase H buffer and the final volume was brought up to 10 μL with nuclease-free water. These samples were mixed thoroughly by pipette and incubated at 37°C for 30 minutes.

At this point, the protocol splits for highly abundant RNA species such as rRNA or snRNA and low abundance RNA species such as mRNA. For highly abundant RNAs, the samples were then incubated at 90°C for 10 minutes to denature the RNase H enzyme and moved to ice. After denaturation, the samples were diluted 1:50 and 5 μL of the dilution was used to make cDNA with SuperScript III (#2563487) and random hexamers. The cDNA was diluted 1:50 and 4 μL of the cDNA dilution was used for RT-qPCR using PowerSybr. For low abundance RNA species, the reaction volume after the 30-minute RNase H cleavage step was brought up to 30 μL with sterile nuclease-free water. The reaction mixture was then transferred to a 1.7 mL microcentrifuge and 30 μL UltraPure™ Phenol:Chloroform:Isoamyl Alcohol (#15593031) was added. After vortexing vigorously, the mixture was spun down for 5 minutes at 12000 g. The upper aqueous phase was transferred to a clean 1.7 mL Eppendorf tube. Cytiva MicroSpin G-50 columns (#27533001) were prepared by loosening the cap, removing the bottom plug, and placing it into a 2 mL collection tube. Excess buffer was removed by spinning for 1 minute at 700 g. The column was then transferred to a clean 1.7 mL Eppendorf tube and the upper phase from the previous extraction step was added to the column. To elute, the tube was spun down at 700 g for 2 minutes.

Primer sequences designed to flank the cleavage sites are listed in the Supplementary Methods. All sequences for experimental and control chimeras are listed in the Supplementary Methods. Data analysis was done using Microsoft Excel and percent methylation was calculated by comparing the cycle numbers of +RNase H enzyme samples to the matching -RNase H samples.

### Luciferase Assay

Luciferase assays were performed using the OZBiosciences Luciferase Assay Kit (#LUC1000). Excess medium was removed from the cell culture plate and the cells were rinsed twice with PBS. 182 μL of 1X cell lysis buffer was added for a 12-well plate and 500 μL of 1X cell lysis buffer was added for a 6-well plate. The plate was incubated at room temperature for 15 minutes before cells were scraped into a 1.7 mL microcentrifuge tube and centrifuged for 5 minutes at 3500 rpm. 50 μL of the supernatant were transferred to a white, clear bottom 96-well plate (#3610) and 100 μL Luciferase Assay Reagent was added to each well. The signal was read using a Tecan SpectraFluor Plus.

### Stable Transfection of Luciferase Chimeras

A firefly Luciferase containing plasmid CMV-LUC2CP/ARE (#62857) and Luciferase containing plasmid with a chimeric intron CMV-LUC2CP/intron/ARE (#62858) were ordered from AddGene. After linearization, the plasmids were run on a 1% agarose gel for quality control to ensure the plasmids were linearized correctly and no contamination was present (not shown). Kill curves with Hygromycin were set up to determine that 0.25mg/mL was the optimal concentration to ensure adequate selection of cells with antibiotic resistance. HEK293T cells were transfected with 1 ug of each plasmid using Lipofectamine. After transfection, cells were grown in 10% FBS DMEM supplemented with 0.25 mg/mL Hygromycin as a selection agent. Selection ran for 10 days before clones were selected and plated into a 96-well plate. After adequate growth occurred, the cells were split and transferred into a 12-well plate to allow for further growth. Selected clones were then tested for luciferase activity to ensure that plasmids were correctly incorporated, and luciferase transcripts and proteins were being expressed.

### snoRNA Abundance

Protocol adapted from Elliot et al^19^. Samples from the transfection experiments were used to make cDNA with SuperScript III. Gene specific primers specific to each snoRNA rather than random hexamers were used to specifically amplify the sequences. 4 μL of the cDNA was used for RT-qPCR using PowerSybr. A universal reverse stem loop primer was used as well as forward primers specific to each snoRNA. All sequences are listed in the Supplementary Methods.

### Statistical Methods

All statistical calculations were performed with GraphPad Prism 10. Hypothesis tests for all Nm-VAQ assays and fold-change assays were calculated using student’s t-test, unpaired, 2-tailed. Hypothesis tests for all Luciferase assays and *Luciferase* mRNA expression assays were calculated using a mixed-effects model and Tukey’s multiple comparisons test with a single pooled variance. All p-values ≤ 0.05 were considered significant.

## Acknowledgements

The authors would like to thank Dr. Brittany Elliot and Dr. Katherine Zhou for their advice and support throughout the process. They would also like to thank Dr. Sheng-Yang He for his feedback in drafting this article.

## Primers and Sequences

### Nm-VAQ RNA-DNA Chimeras

| **Chimera** | **Sequences 5'→3'** |
| --- | --- |
| 18S A159 | dTdTdAdC mCmAmC mAmGmU mUmAmU mCmCmA mAmG |
| 18S U161 | dAdAdTdT mAmCmC mAmCmA mGmUmU mAmUmC mCmA |
| 18S G1328 | dCdCdAdC mCmAmC mCmCmA mCmGmG mAmAmU mCmG |
| U6 C60 | dCdCdAdT mGmCmU mAmAmU mCmUmU mCmUmC mUmG |
| RPL10 G150 | dCdCdAdG mGmUmC mAmAmA mAmAmU mGmCmG mAmA |
| RPL10 G152 | dCdCdCdC mAmGmG mUmCmA mAmAmA mAmUmG mCmG |
| TOP1 U877 | dAdGdCdG mCmUmC mUmUmC mUmUmC mCmCmA mC |
| TOP1 U879 | dAdTdAdG mCmGmC mUmCmU mUmCmU mUmCmC mC |
| *PLXNB2* G498 | dCdAdGdA mGmCmG mCmAmG mAmUmG mCmCmC mU |
| *PLXNB2* G500 | dCdGdCdA mGmAmG mCmGmC mAmGmA mUmGmC mC |
| *Luc* Lys130 | dTdTdGdC mUmCmA mCmGmA mAmUmA mCmGmA mCmG |
| *Luc* Lys130 Control | dTdCdTdT mGmCmU mCmAmC mGmAmA mUmAmC mGmA |

### GSP RT Primers

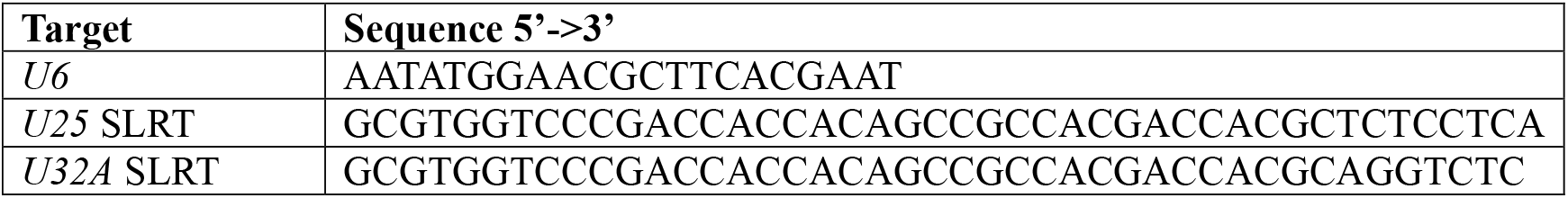

### *In vitro* transcription snoRNA Primers

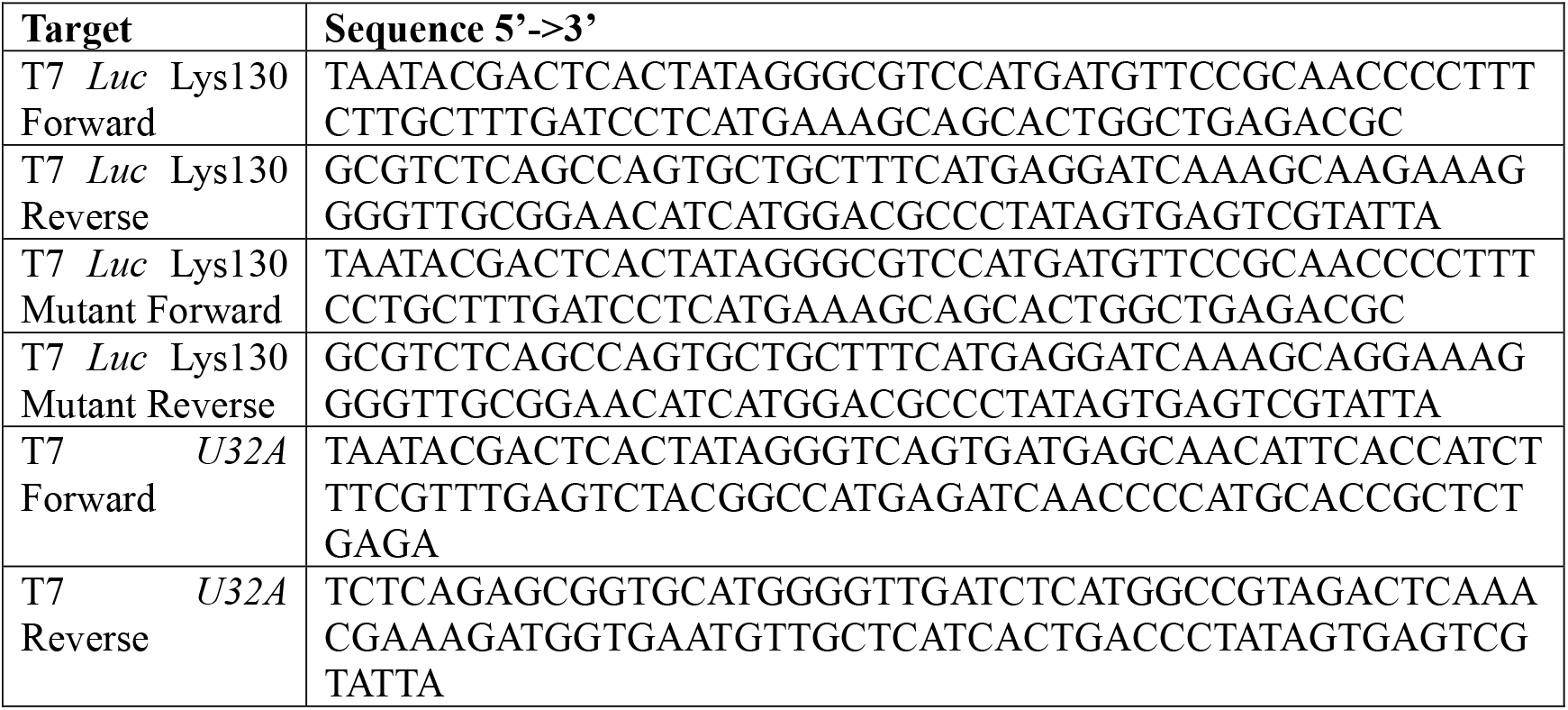

### qPCR Primers

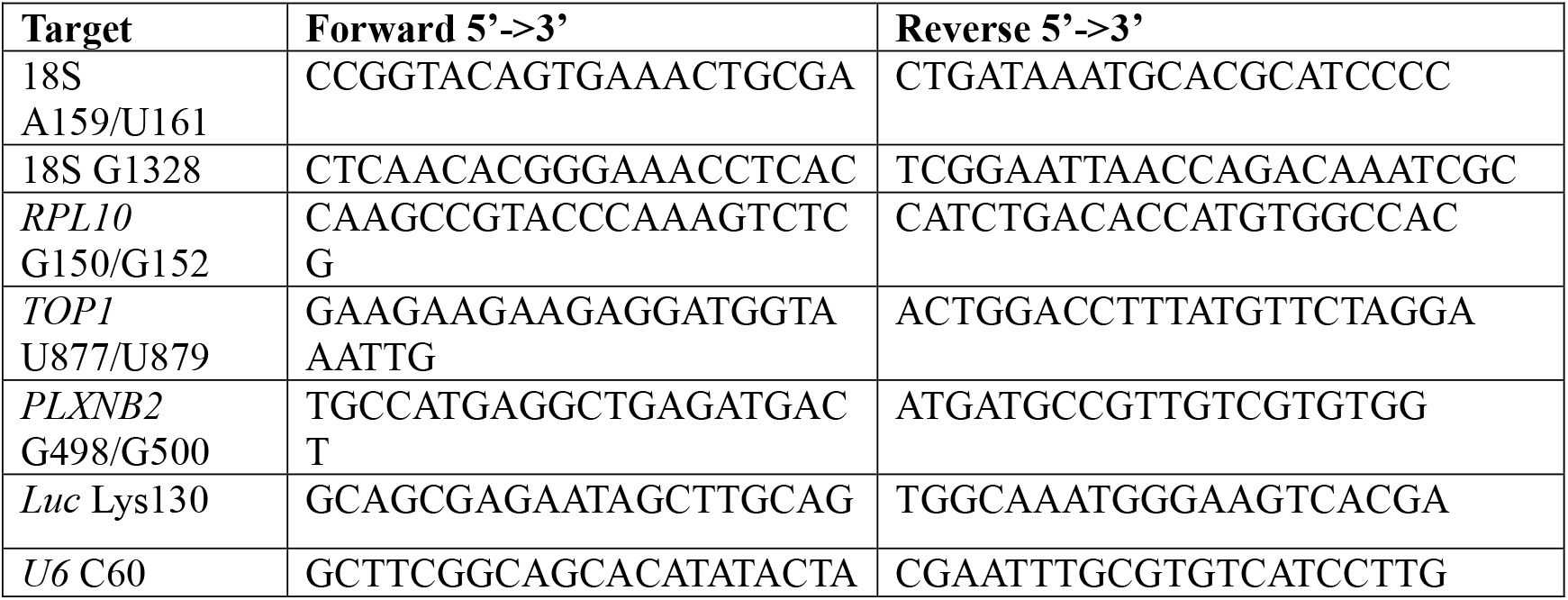
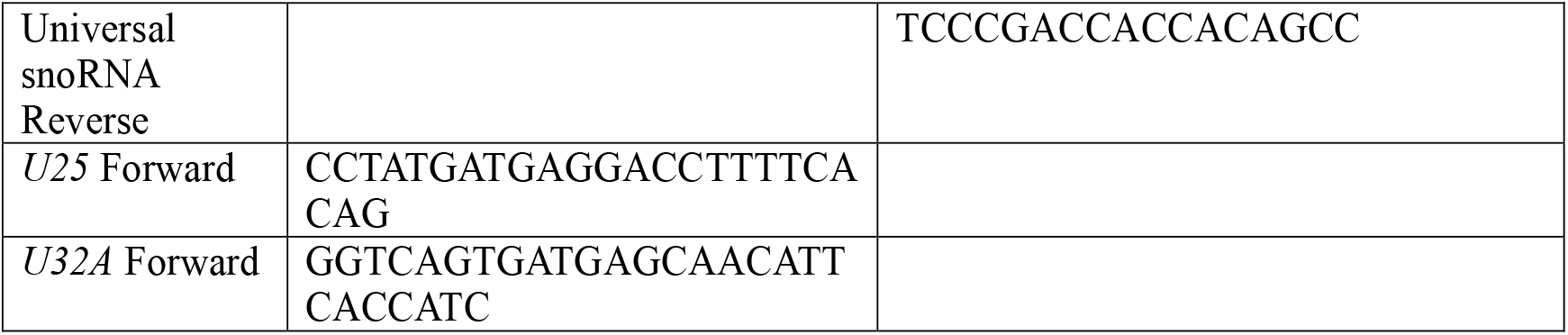

